# Human cytomegalovirus microRNAs target the Wnt signaling protein Dishevelled 3 to promote efficient reactivation in CD34^+^ hematopoietic progenitor cells

**DOI:** 10.64898/2026.10.18.759990

**Authors:** Nicole L Diggins, Luke Slind, Jennifer Mitchell, Andrew H Pham, Devin N Fachko, Yan Chen, Rebekah L Turner, Patrizia Caposio, Rebecca L Skalsky, Meaghan H Hancock

## Abstract

Human cytomegalovirus (HCMV) establishes latency in CD34^+^ hematopoietic progenitor cells (HPCs), yet differentiation of these cells promotes viral gene expression and replication. HCMV must therefore precisely regulate the signaling networks that govern HPC fate to maintain latency while preserving the capacity for reactivation. Virally encoded microRNAs (miRNAs) represent powerful tools used by HCMV to manipulate cellular pathways that are involved in HPC fate decisions. Wnt (Wingless/Integrated)/β-catenin signaling represents one such pathway that plays a critical role in regulating both hematopoiesis and virus replication. Previous work determined that HCMV stimulates β-catenin activity during reactivation in CD34^+^ HPCs. In contrast, here we show that HCMV miRNAs miR-US5-2 and miR-US25-2 downregulate the Wnt pathway adaptor protein Dvl3 during latency and that this downregulation is required for subsequent virus reactivation. These findings support a model of temporally distinct regulation of Wnt pathway components during latency and reactivation.

**IMPORTANCE:** Human cytomegalovirus (HCMV) persists for life by establishing latency in hematopoietic progenitor cells followed by periodic reactivation events. How the virus controls host signaling pathways to influence these events remains incompletely understood. Here, we show that HCMV-encoded microRNAs (miRNAs) downregulate the Wnt signaling adaptor Dvl3 during latency, contrasting with the activation of the Wnt/β-catenin signaling that occurs during reactivation. These findings reveal that HCMV does not uniformly activate or inhibit Wnt signaling but instead regulates this pathway in opposing directions at different stages of infection. This temporal regulation of Wnt signaling highlights how HCMV coordinates viral infection with changes in progenitor cell state and identifies viral miRNAs as important regulators of the balance between latency and reactivation.

## INTRODUCTION

Human cytomegalovirus (HCMV) is a member of the betaherpesvirus family that establishes a lifelong infection due in part to its ability to undergo cycles of latency and reactivation in myeloid-lineage cells. In most immunocompetent hosts, HCMV infection is relatively benign; however, in those undergoing hematopoietic or solid organ transplantation, HCMV reactivation and disseminated infection can cause serious morbidity and even mortality (1). One characterized site of latency in the human host is CD34^+^ hematopoietic progenitor cells (HPCs) (2–4). In the bone marrow, these multipotent HPCs sustain hematopoiesis by responding rapidly to changing physiological demands, while also maintaining the progenitor cell population throughout the lifetime of the organism. These cells respond to extracellular cues and regulate the expression of lineage-specific genes and cell cycle regulators to influence cell development, maintenance and differentiation (5). How HCMV interfaces with and/or integrates into these processes to regulate its own lifecycle during infection of HPCs is only beginning to be appreciated.

The Wnt (Wingless/Integrated)/β-catenin signaling pathway is key developmental pathway that plays an essential role in many fundamental biological processes including embryonic development, organ formation and hematopoiesis and is a frequent target of virally induced dysregulation (6). Secreted Wnt ligands bind to cell surface Frizzled receptors, which, depending on the specific combination of ligand and receptor, can give rise to a variety of downstream signaling cascades (7). Classically, these pathways are referred to as canonical and non-canonical Wnt signaling. β-catenin is the key transcriptional effector of the canonical Wnt signaling pathway. In the absence of Wnt signaling, a multi-protein destruction complex containing GSK3β, CK1, Axin and APC phosphorylates β-catenin, targeting it for ubiquitination and proteasome-dependent degradation (8). Canonical Wnt signaling activates the scaffolding protein Disheveled (Dvl), which promotes recruitment of destruction complex components to the membrane and inhibits β-catenin degradation (9). Stabilized β-catenin accumulates and translocates to the nucleus, where it interacts with TCF/LEF transcription factors to activate Wnt-responsive genes. The Dvl family comprises three ubiquitously expressed paralogs, Dvl1, Dvl2 and Dvl3, which exhibit substantial functional redundancy *in vitro* (10). Non-canonical Wnt signaling also often involves Dvl activation and modulates processes involved in calcium signaling, cell polarity and actin cytoskeletal rearrangements, generally in the absence of β-catenin-mediated transcription (7).

Critically, canonical Wnt signaling functions as a rheostat in hematopoietic progenitors, with different levels of pathway activity associated with distinct cellular outcomes. Relatively low levels of Wnt activity support stem and progenitor cell function, whereas increasing pathway activity can alter lineage commitment and differentiation in a dose-dependent manner. Excessive Wnt signaling, however, impairs hematopoietic function and long-term repopulation capacity (11–13). Thus, precise regulation of Wnt signaling is particularly important in HPCs, where the magnitude, timing and cellular context of pathway activation can influence both the immediate behaviour and subsequent fate of the cell.

Given the central role of Wnt signaling in regulating HPC fate, HCMV regulation of this pathway has the potential to significantly influence the outcome of infection in these cells. Indeed, HCMV modulates Wnt signaling pathway in multiple ways, depending on the context of infection. During lytic infection in fibroblasts, HCMV reduces β-catenin abundance and transcriptional activity and alters components of the Wnt pathway, including LRP6, Axin1 and Tankyrase (14–16). Conversely, the HCMV G protein-coupled receptor US28 can stimulate β-catenin signaling through a non-conventional Rho-ROCK-dependent mechanism (17). Although the functional significance of Wnt pathway regulation during lytic infection remains unclear, a direct role has emerged during HCMV latency and reactivation. The HCMV membrane-associated protein UL8 interacts with Dvl2 and β-catenin to stimulate β-catenin-dependent transcription, and this activity is required for efficient reactivation from latency in CD34^+^ HPCs (18). The requirement for Wnt/β-catenin activity during reactivation raises the question of whether and how this pathway is regulated during latency.

Given the importance of viral microRNAs (miRNAs) in regulating cell signaling during latency, we investigated whether HCMV miRNAs contribute to regulation of Wnt signaling in CD34^+^ HPCs. We and others have shown that HCMV-encoded miRNAs play a pivotal role in modulating signal transduction pathways that can ultimately influence the outcome of HCMV infection in these cells (19–26). Here, we identify the Wnt signaling scaffold protein Dvl3 as an important target of the HCMV miRNAs miR-US5-2 and miR-US25-2. In contrast to the essential role of the paralog Dvl2 in viral reactivation (18), we show that HCMV miRNA-mediated downregulation of Dvl3 during latency is required for efficient reactivation. These findings reveal differential regulation of two Dvl paralogs during HCMV infection of HPCs and support a model in which Wnt pathway components are regulated differently during latency and reactivation.

## RESULTS

### Identification of miR-US5-1 and miR-US5-2 targets using Ago immunoprecipitation

To better understand the role of HCMV miRNAs in the virus lifecycle, it is essential to determine the targeting breadth of each miRNA. In this study, we focused on miR-US5-1 and miR-US5-2 which are encoded adjacent to each other and antisense to the US7 3’UTR, suggesting these miRNAs are derived from the same primary transcript (27). To investigate targets for these two miRNAs, we introduced a lentiviral construct that generates both miR-US5-1 and miR-US5-2 into 293T cells. To map miRNA binding sites, we expanded stably transduced cells, treated cells +/- EGF and performed photoactivatable-ribonucleoside-enhanced crosslinking and immunoprecipitation (PAR-CLIP) analysis (28) using pan-Ago antibodies to isolate RNAs bound by the endogenous Ago2 protein in an unbiased manner (Fig 1A).

**Figure 1.**
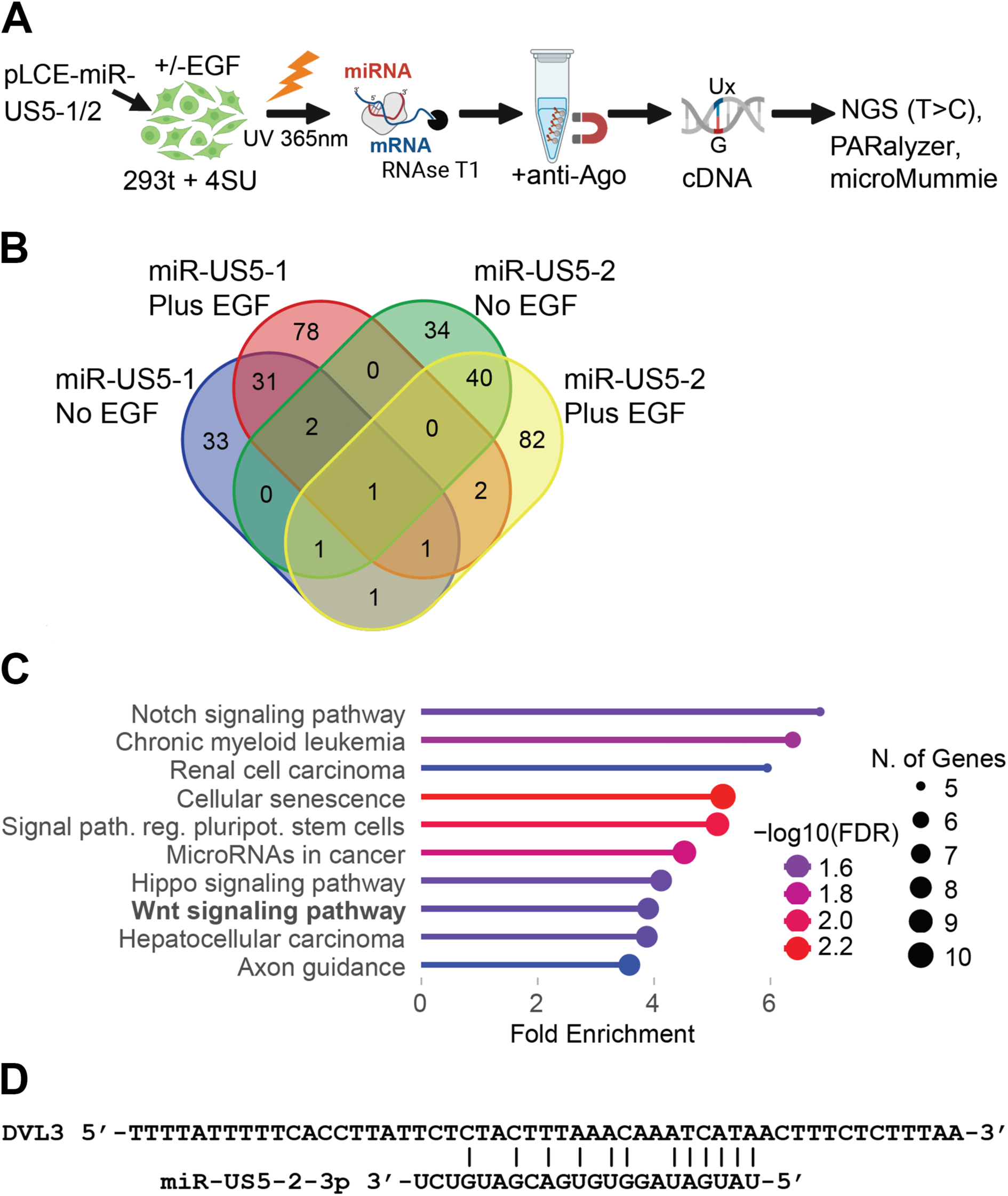
Identification of HCMV miR-US5-1 and miR-US5-2 targets. A) Experimental design for PAR-CLIP analysis. HEK293 cells infected with lentivirus expressing miR-US5-1 and miR-US5-2 and were treated with or without EGF for 22hr as well as 4SU. Ago-associated mRNAs were isolated, sequenced and analyzed as previously described. B) Venn diagram shows overlap of the number of human protein-coding genes targeted by each miRNA. C) ShinyGO 0.77 pathway analysis of miR-US5-1 and miR-US5-2 targets. Shown are pathways from the KEGG database. FDR cutoff >=0.1, pathway size >=10. D) Region within the DVL3 3’UTR captured by PAR-CLIP with the miR-US5-2-3p seed match site.

Following immuno-purification, Ago-bound RNAs were converted to cDNA and sequenced. PAR-CLIP reads were then analyzed by PARalyzer (29) and microMUMMIE (30) to identify Ago/miRNA binding sites with T-to-C conversions. To discern the HCMV miRNA binding sites, CLIP’ed sites within 3’UTRs of human protein coding transcripts were scanned for canonical seed matches (>=6mer2-7). In total, we defined 186 3’UTR target sites for miR-US5-1-3p, 200 target sites for miR-US5-2-3p, and 10 target sites for miR-US5-2-5p (Table S1). Of these, 110 genes were previously identified as miR-US5-1 and miR-US5-2 targets by Kim *et al* in Ago-CLIP-seq studies (31), while 196 genes were novel HCMV miRNA targets identified in the current study (Fig 1B). Although relatively few targets of these miRNAs have been functionally validated, (22, 27, 31, 32), the analysis recovered CHUK/IKKα, a previously validated miR-US5-1 target (32), supporting the ability of this approach to identify biologically relevant targets.

To understand the pathways targeted by miR-US5-1 and miR-US5-2, we used the KEGG pathway analysis tool (33) to analyze the HCMV miR-US5-1/5-2 target list generated by PAR-CLIP analysis (Fig 1C). Amongst the pathways enriched for HCMV miRNA targets was R-HSA-201681, “TCF dependent signaling in response to WNT” and included targets such as CREBBP, PSMD12, APC, PSMB2, TNKS, CTBP1, DVL3, FZD8 and YWHAZ. Of the members of this signaling pathway, our computational analysis identified Dvl3 as a target of miR-US5-2 (Fig 1D). Given our previous studies into the role of Wnt signaling in HCMV reactivation (18), we chose to follow up on the role of HCMV miRNAs in regulating this signaling pathway.

### HCMV miR-US5-2 and miR-US25-2 suppress β-catenin activation

Because elevated levels of Wnt signaling can alter HPC differentiation and impair progenitor cell function (12, 13), we hypothesized that HCMV suppresses Wnt activity during latent infection. To determine if HCMV miRNAs can affect signaling through the canonical Wnt pathway, we tested the ability of a subset of HCMV miRNAs to inhibit Wnt3a-mediated activation of a TCF/LEF luciferase reporter in HEK293 cells. Figure 2A shows that expression of HCMV miR-US5-2 and miR-US25-2 was able to inhibit Wnt3a-mediated luciferase expression, while miR-US4-1 had no effect. We utilized a control reporter, which contains mutations in the TCF/LEF binding sites, to verify the specificity of the effects of miR-US5-2 and miR-US25-2 (Fig 2B). To better understand where in the Wnt signaling pathway the HCMV miRNAs were acting, we also stimulated transfected cells with LiCl, a GSK3β inhibitor that influences the composition of the destruction complex (34, 35). We found that LiCl was able to enhance luciferase expression regardless of miRNA expression, indicating that the miRNAs function upstream of the destruction complex (Fig 2C). We also tested whether the miRNAs, alone and in combination, or a Dvl3 siRNA could affect active (non-phosphorylated) β-catenin levels in human fibroblasts. As shown in Figure 2D-F, transfection of miRNA mimics or Dvl3 siRNA in normal human dermal fibroblasts (NHDF) significantly reduced the level of active β-catenin, while having no effects on total β-catenin levels. Thus, these findings suggest that the miRNAs act upstream of GSK3β-dependent β-catenin stabilization.

**Figure 2.**
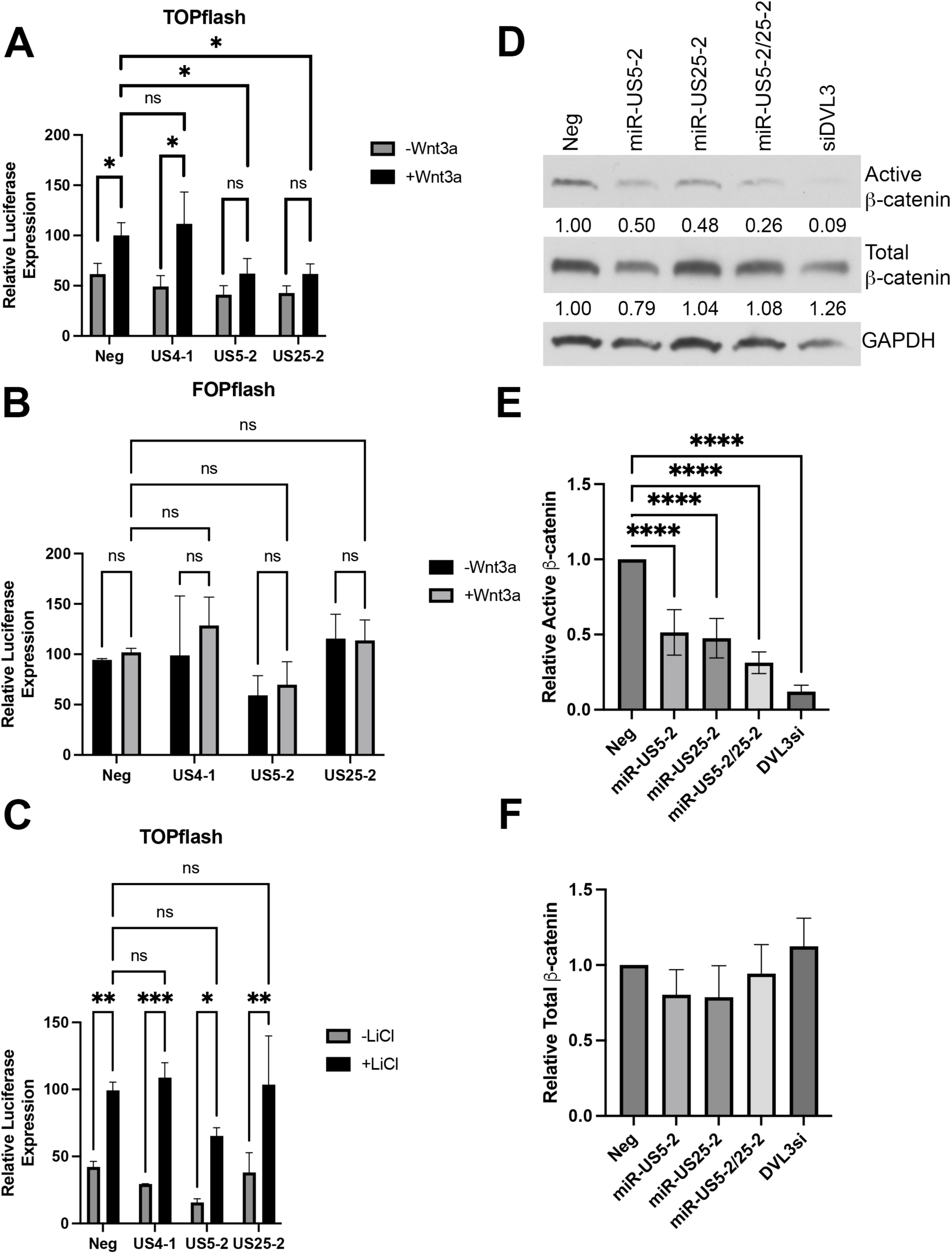
HCMV miRNAs miR-US5-2 and miR-US25-2 inhibit β-catenin signaling. (A-C) Tcf/Lef (TOPflash) reporter plasmid (A,C) or a reporter plasmid containing mutated TCF binding sites (FOPflash) (B) was co-transfected with negative control or miRNA mimics in HEK293T cells. Cells were serum starved for 16hr, treated +/- 150ng/ml Wnt3a (A, B) or 25mM LiCl (C) for 8hr, then assessed for luciferase activity. (D) Normal human dermal fibroblast (NHDF) cells were transfected with miRNA mimics, Dvl3 siRNA or negative control mimic. Cells were lysed 72hr later and subjected to western blot analysis. Quantification of active or total β-catenin levels are normalized to GAPDH and Neg. (E, F) Quantification of active (E) or total (F) β-catenin levels from three separate experiments. *p<0.05, **p<0.01, ***p<0.001 by two-way ANOVA (A, C), ****p<0.0001 by one-way ANOVA (E).

### HCMV miR-US5-2 and miR-US25-2 downregulate Dvl3 protein levels

That ability of miR-US5-2 to restrict Wnt/β-catenin signaling is consistent with the identification of a miR-US5-2 binding site within the Dvl3 3’UTR by PAR-CLIP. We have previously shown that multiple HCMV miRNAs can target the same transcript to enhance repression of protein expression (20, 22, 27, 32, 36). Given that miR-US25-2 also reduced Wnt signaling, we asked whether this miRNA similarly regulates Dvl3 expression. We cloned the 3’UTR of Dvl3 into a luciferase reporter construct and assessed the ability of negative control or miR-US25-2 mimics to regulate luciferase expression. As shown in Figure 3A, miR-US25-2 significantly reduced luciferase expression regulated by the Dvl3 3’ UTR. We next tested whether miR-US5-2 and miR-US25-2 could regulate endogenous Dvl3 protein levels. Human fibroblasts were transfected with negative control mimic, miR-US5-2 and miR-US25-2, alone or in combination, as well as a Dvl3 siRNA. As shown in Figure 3B-C, the Dvl3 siRNA and each miRNA alone or in combination reduces Dvl3 protein levels. These data indicate that miR-US5-2 and miR-US25-2 regulate Wnt/β-catenin signaling (Fig 2), at least in part through regulation of Dvl3 protein expression.

**Figure 3.**
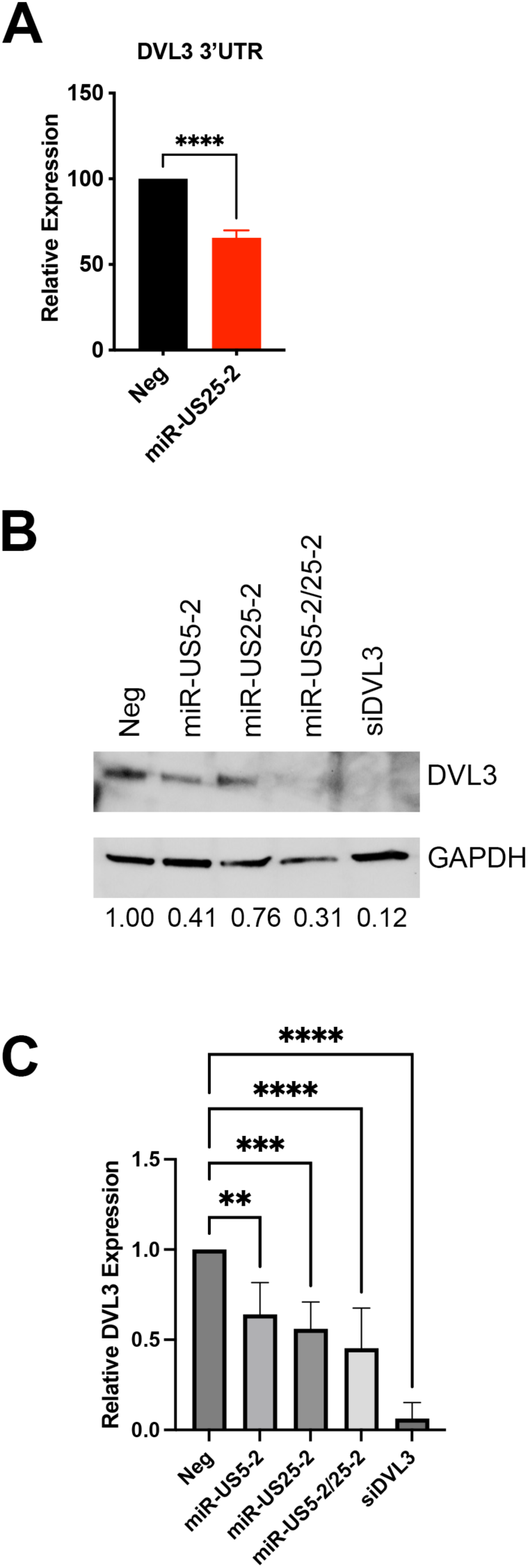
HCMV miR-US5-2 and miR-US25-2 target Dvl3. (A) A dual luciferase reporter for the DVL3 3’UTR was transfected into HEK293T cells along with miRNA mimics. After 24hr, cells were lysed and assessed for luciferase activity. (B) NHDFs were transfected with miRNA mimics, Dvl3 siRNA or negative control mimic. Cells were lysed 72hr later and subjected to western blot analysis. Quantification of Dvl3 protein levels were normalized to GAPDH and Neg. (C) Quantification of (B) from three separate experiments. **p<0.01, ***, p<0.001, ****p<0.0001 by student’s t-test (A) or one-way ANOVA (C).

### miR-US5-2 and miR-US25-2 target Dvl3 during infection

To investigate the importance of Dvl3 regulation in the context of HCMV infection, we generated a recombinant TB40/E virus where both miR-US5-2 and miR-US25-2 miRNA hairpins were mutated. Additionally, we generated a recombinant construct where miR-US5-2 was mutated and an shRNA targeting Dvl3 replaced the miR-US25-2 hairpin, allowing the Dvl3 shRNA to be expressed with the same kinetics as the viral miRNA (Fig 4A). Replication of the recombinant viruses was similar to WT as assessed by both single and multi-step growth kinetics (Fig. S1), indicating that loss of the miRNAs and Dvl3 knockdown did not affect lytic replication. To assess miRNA regulation of Dvl3 protein expression during infection, human fibroblasts were infected with WT or the above recombinant viruses at a multiplicity of infection (MOI) of 3 and protein lysates were harvested 72 and 96 hours later. As shown in Figure 4B-C, cells infected with WT virus showed reduced Dvl3 protein levels compared to mock-infected cells at both timepoints, while in ΔmiR-US5-2/25-2-infected cells, Dvl3 levels were significantly higher than in WT-infected cells at 96hpi. Conversely, the ΔmiR-US5-2/25-2/DVL3 shRNA virus infection resulted in decreased Dvl3 expression compared to parental ΔmiR-US5-2/25-2 infected cells and to levels similar to WT infection, indicating that the Dvl3 shRNA was able to complement the loss of miR-US5-2 and miR-US25-2 in regulation of Dvl3 protein levels. We also assessed Dvl3 protein levels in CD34^+^ HPCs after 14 days of infection and observed a similar trend, where infection of HPCs with the ΔmiR-US5-2/25-2 recombinant virus resulted in enhanced Dvl3 protein levels compared to WT-infected cells, while infection with the ΔmiR-US5-2/25-2/DVL3 shRNA virus resulted in reduced Dvl3 protein detection (Fig. 4D). These data indicate that miR-US5-2 and miR-US25-2 regulate DVL3 protein levels in fibroblasts and CD34^+^ HPCs, and this regulation can be manipulated through expression of an shRNA targeting DVL3 from the viral genome.

**Figure 4.**
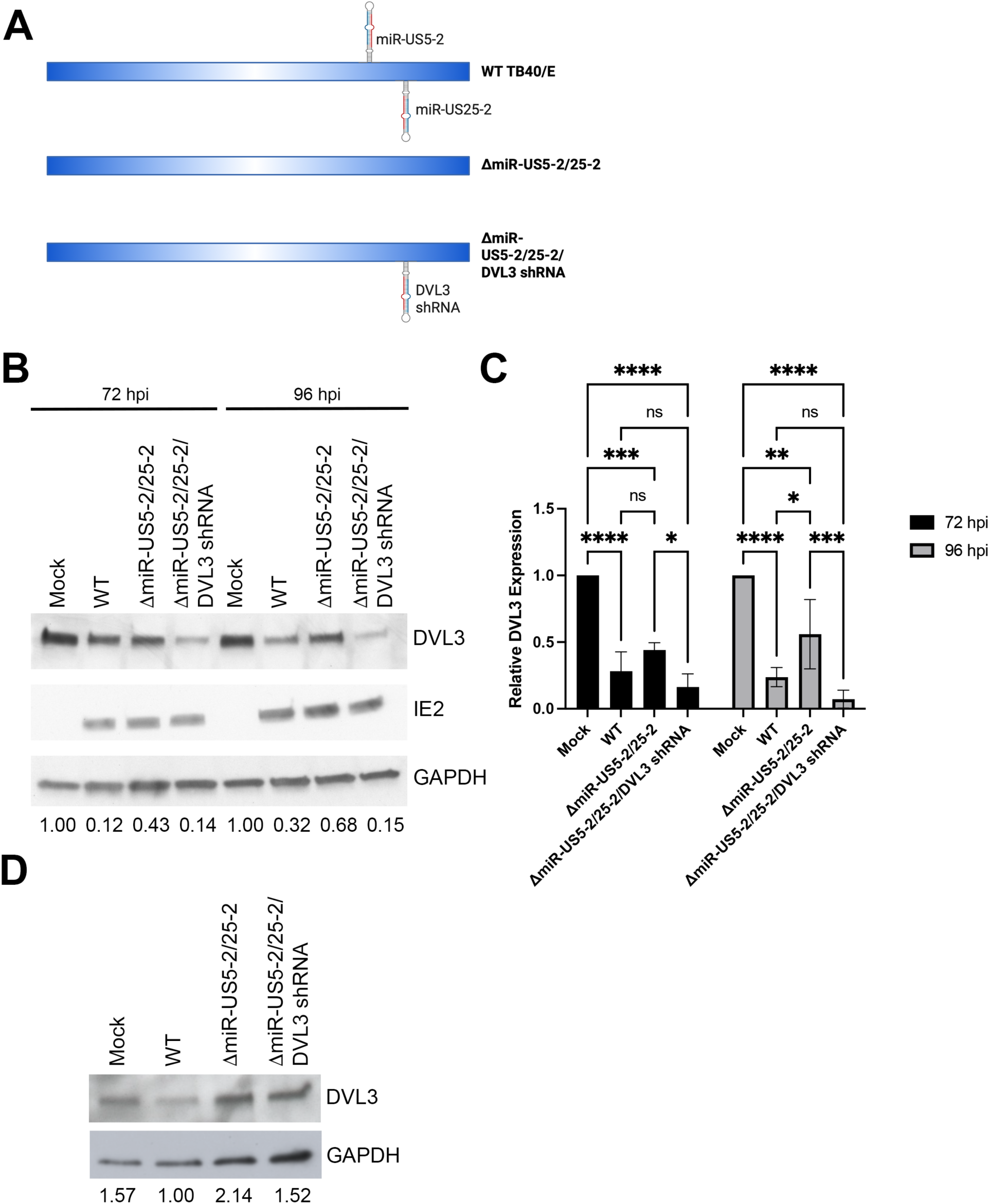
miR-US5-2 and miR-US25-2 modulate Dvl3 during HCMV infection. (A) Schematic of mutant viruses. (B) NHDFs were infected at an MOI of 3 with WT HCMV (TB40/E-GFP), a mutant lacking miR-US5-2 and miR-US25-2 expression (ΔmiR-US5-2/25-2), a mutant lacking miR-US5-2 and miR-US25-2 but expressing an shRNA targeting DVL3 (ΔmiR-US5-2/25-2/DVL3 shRNA), or uninfected (Mock). After 72 and 96hr, cells were lysed and subjected to western blot analysis. Quantification of DVL3 levels were normalized to GAPDH and Mock. (C) Quantification of (B) from three separate experiments. (D) hESC-derived CD34^+^ HPCs were infected at an MOI of 2 with WT, ΔmiR-US5-2/25-2, or ΔmiR-US5-2/25-2/DVL3 shRNA, or Mock. After 48hr, cells were sorted by FACS for viable, CD34^+^, GFP^+^ cells. HPCs were cultured in SFEM hematopoietic cell media overnight and then lysates were harvested and immunoblotted for DVL3 and GAPDH. Quantification of DVL3 levels were normalized to GAPDH and WT. *p<0.05, **p<0.01, ***p<0.001, ****p<0.0001 by one-way ANOVA.

### Regulation of Dvl3 is necessary for efficient reactivation from latency

Since our previous work highlighted the important role of Wnt/β-catenin signaling in HCMV reactivation from latency (18), we next sought to determine the role of Dvl3 regulation in the context of HCMV infection of CD34^+^ HPCs. We infected human embryonic stem cell (hESC)-derived CD34^+^ HPCs with WT and recombinant viruses using our standard latency and reactivation protocol (37, 38). In brief, hESCs were differentiated into HPCs and infected with WT, ΔmiR-US5-2/25-2, or ΔmiR-US5-2/25-2/DVL3 shRNA viruses at an MOI of 2. At 48 hpi, cells were sorted by FACS for viable, CD34^+^, GFP^+^ cells and then seeded into long-term bone marrow culture (LTBMC) as previously described (37, 38). After 12 days in latency culture, cells were plated onto permissive fibroblasts in cytokine-rich media in an extreme limiting dilution assay (ELDA;(39)), and the frequency of infectious center production was scored by GFP^+^ wells after three weeks of co-culture. An equivalent number of cells were lysed and plated in the ELDA, serving as a pre-reactivation control. As shown in Figure 5A and S2, virus lacking expression of miR-US5-2 and miR-US25-2 did not reactivate from latency as effectively as WT virus, demonstrating that these viral miRNAs promote efficient reactivation. Importantly, expression of a Dvl3 shRNA was able to complement the reactivation defect of the miRNA mutant virus, indicating that targeting Dvl3 for downregulation is a major function of miR-US5-2 and miR-US25-2 in CD34^+^ HPCs. Viral DNA levels were similar in all conditions at the time of reactivation (Fig. 5B and S3), indicating that the defect of the miRNA mutant virus was due to an inability to re-initiate the virus replication cycle rather than through loss of viral genomes or genome-containing cells during latent infection.

**Figure 5.**
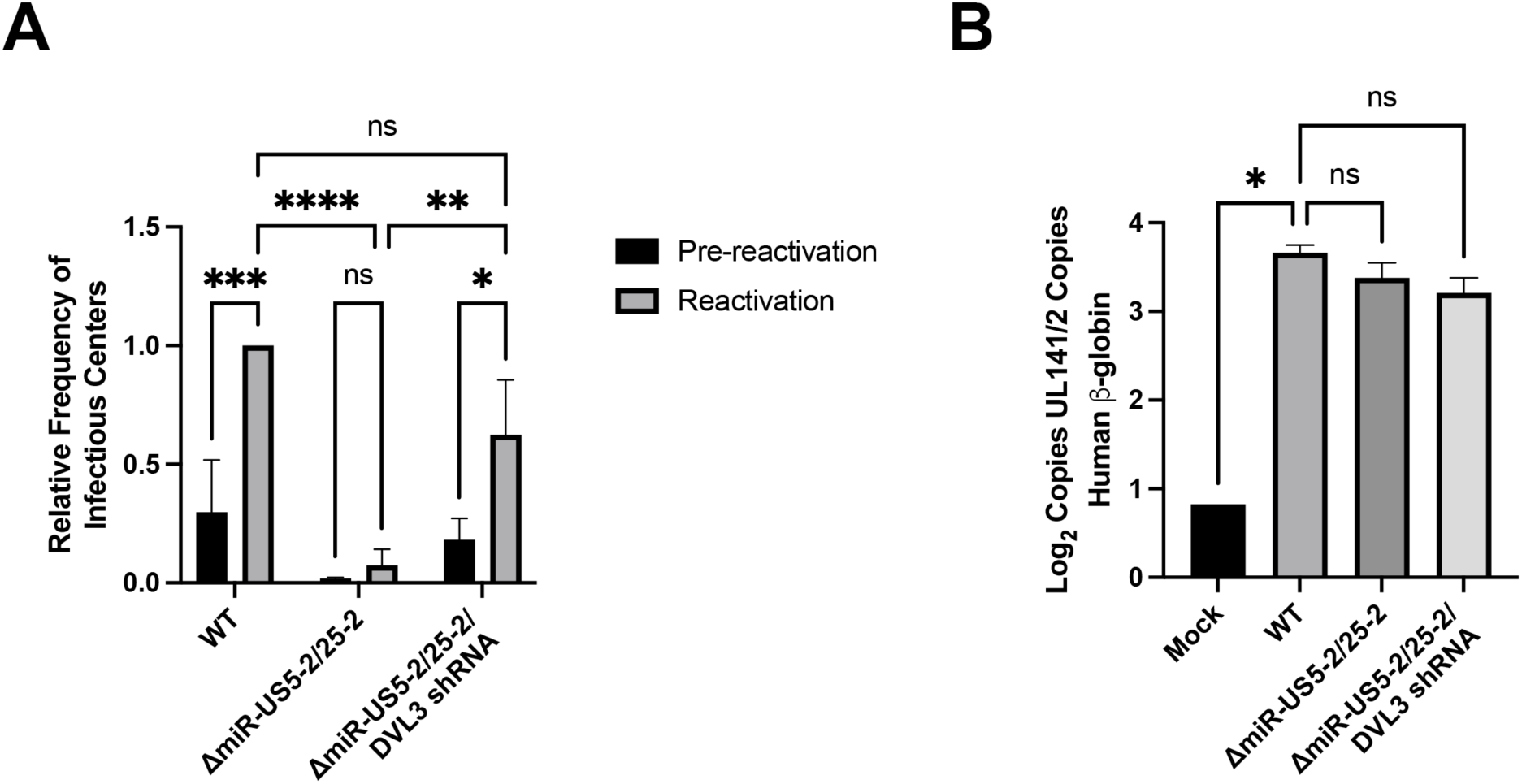
Targeting of Dvl3 by miR-US5-2 and miR-US25-2 is essential for efficient reactivation. hESC-derived CD34^+^HPCs were infected at an MOI of 2 with WT, ΔmiR-US5-2/25-2, ΔmiR-US5-2/25-2/DVL3 shRNA, or Mock. After 48hr, cells were sorted by FACS for viable, CD34^+^, GFP^+^ cells. Infected HPCs were maintained in LTBMC culture medium in transwells over stromal cells for 12 days to establish latency. (A) Latently-infected cells were co-cultured with NHDFs in cytokine-rich media in an extreme limiting dilution assay (ELDA) to measure viral reactivation. An equal number of cells were mechanically disrupted and seeded in parallel to measure infectious virus present in the latency culture (pre-reactivation). At 21 days post-plating, the number of GFP^+^ wells were scored and the relative frequency of infectious center production compared to WT-infected cells from three separate experiments. (C) Total genomic DNA was isolated from HPCs at 14 dpi (latency day 12), and quantitative real-time PCR was used to quantify the ratio of viral genomes (copies of HCMV UL141) to cellular genomes (per two copies of human β-globin). Error bars represent standard deviation from triplicate samples. Data shown are representative of three independent experiments. *p<0.05, **p<0.01, ***p<0.001, ****p<0.0001 by two-way ANOVA (A) or one-way ANOVA (B).

### Regulation of Dvl3 during latent infection is necessary for subsequent reactivation

The requirement for miRNA-mediated suppression of Dvl3 for efficient reactivation was particularly intriguing given our previous finding that HCMV UL8 engages the Dvl paralog Dvl2 to promote β-catenin-mediated signaling and reactivation (18). Together, these observations suggest that HCMV differentially regulates Dvl paralogs in latency versus reactivation, potentially reflecting distinct functions of Dvl2 and Dvl3 in HPCs and/or a switch from restriction of Wnt signaling to its activation during reactivation. Because miR-US25-2 is continuously expressed in HPCs (24), our approach of encoding a Dvl3-targeting shRNA within this region of the HCMV genome does not restrict Dvl3 downregulation to latency but also maintains its suppression during reactivation. Thus, we cannot separate the role of Dvl3 in latency from a role in reactivation using this recombinant virus design. As one approach to determine whether HCMV miRNA targeting of Dvl3 is essential during the latent phase of infection, we turned to small peptide inhibitors of Dvl proteins that we could add at discrete phases of infection. Pen-N3 is a cell-permeable peptide that binds to the PDZ domain of Dvl and inhibits Dvl-dependent canonical Wnt signaling, while the Pen peptide is its inactive analog (40). When we added Pen-N3 to HCMV latency cultures, replenishing the drug at 7dpi, we did not see any significant effect on the ability of WT virus to reactivate compared to cells treated with Pen (Fig. 6). However, we did note a significant increase in the ability of the ΔmiR-US5-2/25-2 mutant to reactivate in the presence of Pen-N3, supporting an important role for miRNA targeting of Dvl proteins during latency in the subsequent ability of the virus to reactivate. Moreover, Pen-N3 had no significant effect on the ability of the Dvl3 shRNA virus to reactivate, suggesting that the effect of the peptide inhibitor is mainly mediated through targeting Dvl3. Together, these data indicate that HCMV uses miR-US5-2 and miR-US25-2 to target Dvl3 for downregulation during latent infection, which is critical to the ability of the virus to reactivate efficiently from latency in CD34^+^ HPCs.

**Figure 6.**
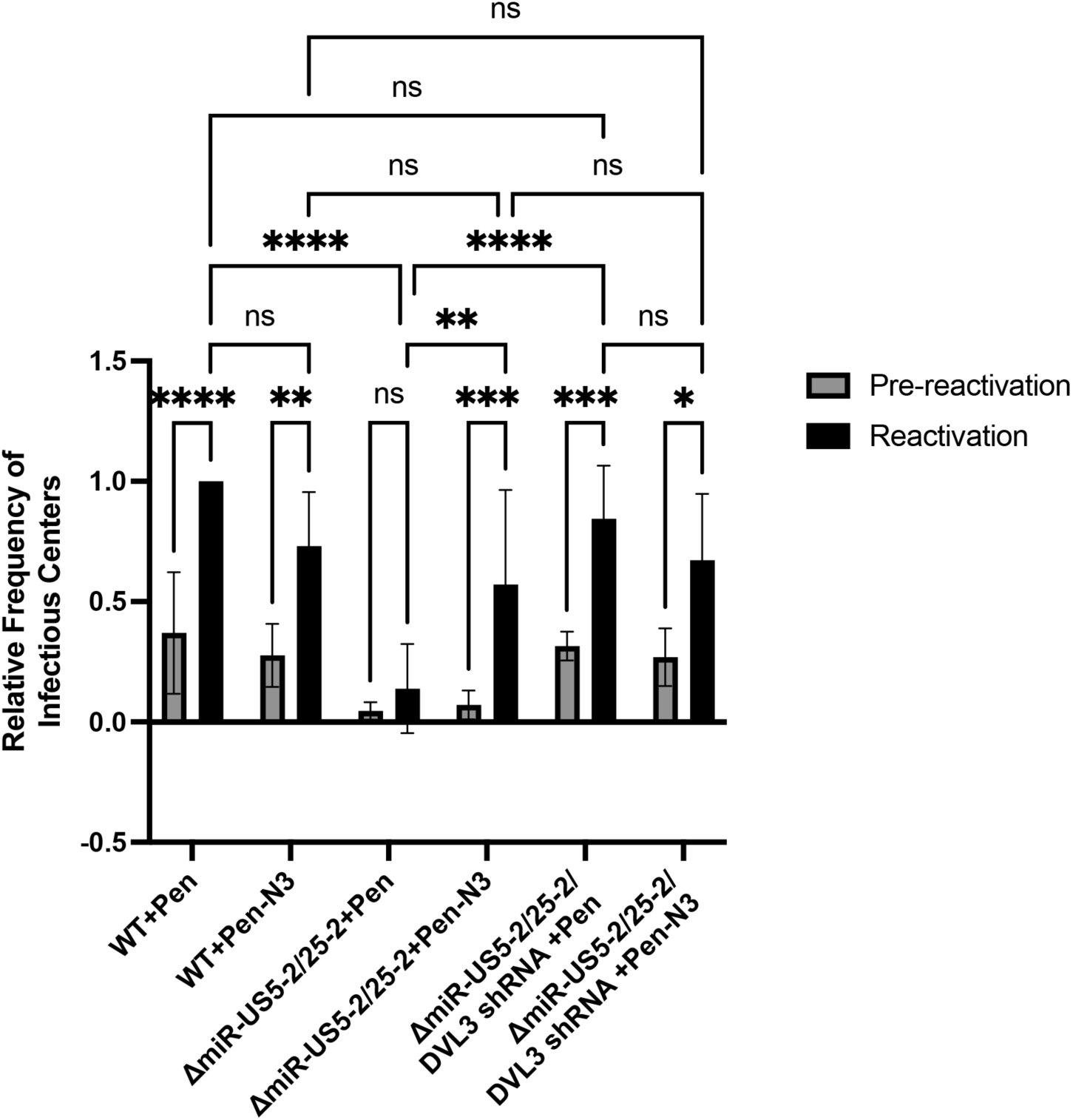
Dvl3 targeting by miR-US5-2 and miR-US25-2 during latency is necessary for reactivation. hESC-derived CD34^+^ HPCs were infected at an MOI of 2 with WT, ΔmiR-US5-2/25-2, or ΔmiR-US5-2/25-2/DVL3 shRNA. After 48hr, cells were sorted by FACS for viable, CD34^+^, GFP^+^ cells. Infected HPCs were maintained in LTBMC culture medium in the presence of 10uM DVL inhibitor (Pen-N3) or control peptide (Pen) and latency and reactivation assays were performed as described in Figure 5. Relative frequency of infectious centers is representative of seven separate experiments. *p<0.05, **p<0.01, ***p<0.001, ****p<0.0001 by two-way ANOVA.

## DISCUSSION

The work presented here expands our understanding of how HCMV manipulates Wnt/β-catenin signaling during infection of CD34^+^ HPCs. We demonstrate that HCMV miRNAs miR-US5-2 and miR-US25-2 target the Wnt scaffolding protein Dvl3 and that this regulation is required for efficient viral reactivation. Together with our previous finding that UL8 promotes Wnt/β-catenin-mediated signaling during reactivation (18), these results support a model in which HCMV differentially regulates Wnt pathway components during latency and reactivation. Thus, rather than simply activating or suppressing Wnt/β-catenin signaling, HCMV exerts temporal control over this pathway, highlighting the importance of precisely regulating Wnt signaling during the transition between latency and reactivation.

Unbiased Ago-IP-based analysis of the HCMV miRNAs miR-US5-1 and miR-US5-2 identified 396 targets, including newly identified and previously reported miRNA targets (31, 32). Pathway enrichment analysis highlighted the Wnt/β-catenin signaling pathway, with miR-US5-2 identified as directly targeting the 3’ UTR of the Wnt scaffolding protein Dvl3. We further determined that miR-US25-2 suppresses Wnt/β-catenin signaling, and downregulates Dvl3, adding to the growing list of cellular proteins and signaling pathways that are regulated by multiple HCMV miRNAs (20, 22, 27, 32). Several lines of evidence indicate that miR-US5-2 and miR-US25-2 regulate Wnt/β-catenin signaling upstream of the β-catenin destruction complex. Neither miRNA affected LiCl-induced β-catenin signaling, which bypasses receptor-proximal events through inhibition of GSK3β (34, 35). Furthermore, the miRNAs reduced levels of active β-catenin without altering total β-catenin abundance, consistent with regulation of β-catenin activity rather than expression. Together, these findings indicate that the effects of miR-US5-2 and miR-US25-2 are on an upstream step in the Wnt pathway and support Dvl3 as a target through which HCMV miRNAs restrict canonical Wnt signaling.

Critically, we demonstrate that miR-US5-2 and miR-US25-2 regulate Dvl3 expression in HPCs and that this regulation is essential for efficient viral reactivation. Because both miRNAs have additional predicted and validated targets (23, 27, 31, 36, 41, 42), we sought to determine whether dysregulation of Dvl3 specifically accounts for the reactivation defect of the miRNA mutant. Expression of a Dvl3-targeting shRNAs from the ΔmiR-US5-2/miR-US25-2 virus restored reactivation to WT levels, demonstrating that Dvl3 downregulation is sufficient to complement the loss of both viral miRNAs. As an orthogonal approach, treatment with the pan-Dvl inhibitory peptide Pen-N3 (40) enhanced reactivation of the miRNA mutant but provided no additional benefit to the ΔmiR-US5-2/miR-US25-2/DVL3 shRNA virus. Together, genetic complementation and pharmacological inhibition identify Dvl3 as a relevant target of miR-US5-2 and miR-US25-2 during latency and demonstrate that miRNA-mediated regulation of Dvl3 is necessary for efficient reactivation.

Our finding that miRNA-mediated downregulation of Dvl3 during latency is necessary for efficient reactivation, while dispensable for maintenance of the viral genome, suggests that maintenance of the latent viral genome and preservation of a cellular state competent for reactivation may represent distinct requirements of latent infection. This distinction may be particularly important in highly dynamic cells such as HPCs, in which changes in signaling can alter differentiation and self-renewal and thus the outcome of HCMV infection. The consequences of Wnt signaling in hematopoietic progenitors are highly dependent on signal strength, timing and cellular context (11–13), suggesting that precise control, rather than simply activation or inhibition, is necessary to maintain progenitor cell function. Loss of miR-US5-2- and miR-US25-2-mediated regulation during latency could therefore alter the developmental trajectory or functional state of infected HPCs, ultimately generating cells that are less competent to support reactivation. We speculate that successful reactivation depends not only on the signals encountered at the time of reactivation but also on the cellular state established during latency. This concept is consistent with US28 as an HCMV protein that act during latency to modify HPC cell state and signaling yet is ultimately required for efficient reactivation (43). HCMV miRNA regulation of progenitor cell signaling during latency may therefore preserve a cellular environment capable of responding appropriately to subsequent reactivation signals.

As miR-US5-2 and miR-US25-2 are expressed throughout both latency and reactivation, we propose a model (Fig 7) whereby miRNA-mediated downregulation of Dvl3 during latency is critical for maintaining the infected progenitor cell in a state capable of supporting reactivation. At the time of reactivation, the viral miRNAs continue to target Dvl3, yet newly expressed HCMV UL8 sequesters Dvl2 and β-catenin to promote β-catenin-mediated transcription, bypassing the suppressive effects of the viral miRNAs on the Wnt signaling pathway. The differential regulation of Dvl paralogs by HCMV – targeting one for downregulation while sequestering another – raises important questions about their paralog-specific functions, which remain poorly defined in human hematopoietic progenitors. Studies in cultured cells suggest substantial functional overlap among the Dvl paralogs, as individual Dvl proteins can support multiple canonical Wnt signaling responses in the absence of the other two family members (44). However, differences in signaling capacity among the individual paralogs, together with distinct phenotypes observed in Dvl knockout mice (45), indicate that Dvl1, Dvl2 and Dvl3 also have context-dependent, non-redundant functions *in vivo*. This context-dependent redundancy may be particularly relevant in hematopoietic progenitors, where regulation of individual Dvl paralogs by HCMV could have consequences beyond their shared roles in canonical Wnt signaling. Thus, several outstanding questions remain to be addressed regarding this model: (1) Is Dvl3 the only paralog involved in Wnt signaling during HCMV latency? If not, does the virus have additional mechanisms to limit signaling mediated by other Dvl paralogs? (2) Is regulation of Dvl3 necessary to limit β-catenin signaling or a non-canonical function of Dvl proteins? (3) Is blocking Dvl3-mediated signaling also essential at the time of reactivation? Unfortunately, no paralog-specific peptide or small molecule inhibitors exist to determine paralog-specific functions during HCMV infection.

**Figure 7.**
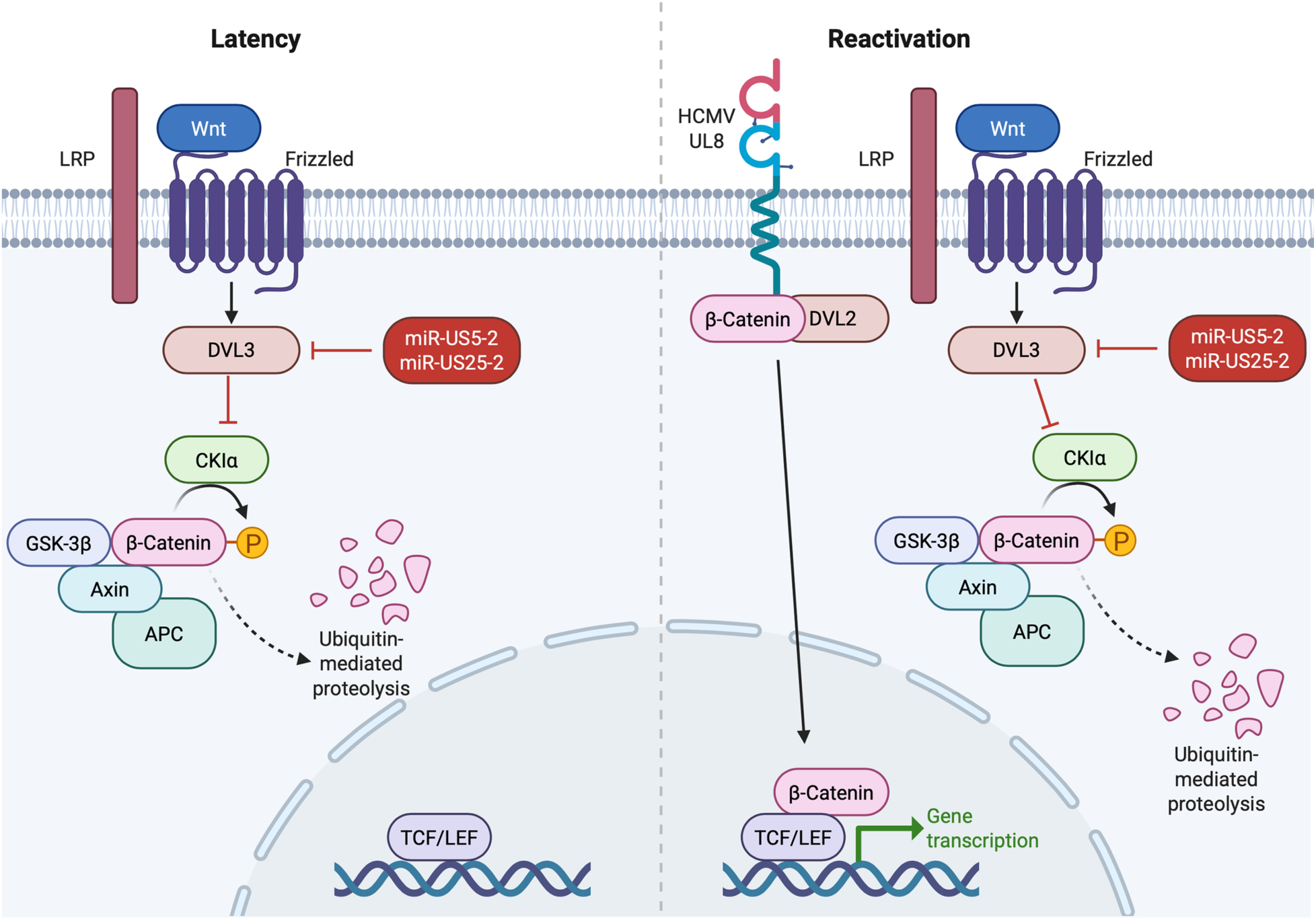
Model of Wnt regulation in latency and reactivation. Left, Wnt signaling recruits Dvl3 to block the destruction complex, thereby turning on Wnt signaling. HCMV miR-US5-2 and miR-US25-2 target Dvl3 to suppress Wnt signaling during latency. Right, UL8 expression during reactivation directly recruits Dvl2 and β-catenin, bypassing the miRNA-mediated block in Wnt signaling and stimulating viral reactivation. Figure generated using BioRender.

The opposing nature of the regulation of Wnt/β-catenin signaling in latency versus reactivation highlights how HCMV has evolved both to reinforce the progenitor phenotype of infected cells and drive their differentiation when appropriate conditions have been met, here utilizing both viral proteins and non-coding RNAs. The differential regulation of Wnt signaling across latency and reactivation becomes particularly interesting when considered together with HCMV-mediated control of Notch signaling (21, 46, 47). Wnt and Notch can exert opposing influences on hematopoietic differentiation, with elevated Notch signaling restraining myeloid differentiation (48, 49), whereas increased Wnt signaling in some contexts promotes myeloid-biased progenitor expansion and myeloid differentiation (50). We have previously shown that HCMV UL8 and the viral miRNA miR-UL36 suppress Notch signaling at the time of reactivation (21). Together with our findings that HCMV miRNAs suppress Wnt signaling during latency, whereas UL8 promotes Wnt signaling during reactivation, these observations suggest a coordinated viral strategy that shifts the balance between two opposing developmental pathways across latency and reactivation. Thus, rather than targeting Wnt and Notch independently, HCMV appears to integrate their regulation in a temporally controlled manner to reshape signaling networks that govern HPC fate.

## Materials and Methods

### Cells and media

Feeder-free hESCs are from WiCell (WA01-H1, hPCSC Reg identifier (ID) WAe001-A, NIH approval no. NIHhESC-10-0043). Cells were thawed and plated on Matrigel-coated six-well plates in complete mTeSR1 media (Stem Cell Technologies). CD34^+^ HPCs were differentiated using a commercial feeder-free hematopoietic differentiation kit (STEMdiff Hematopoietic Kit, Stem Cell Technologies). HEK293 and adult normal human dermal fibroblasts (NHDF) were obtained from ATCC and cultured in Dulbecco’s modified Eagle’s medium (DMEM) supplemented with 10% heat-inactivated fetal bovine serum (FBS; Hyclone), 100 units/ml penicillin, 100 μg/ml streptomycin, and 100 μg/ml glutamine (Thermofisher). M2-10B4 and S1/S1 stromal cells were obtained from Stem Cell Technologies and maintained in DMEM with 10% FBS and penicillin, streptomycin, and glutamine as previously described. All cells were maintained at 37°C and 5% CO_2_.

### Viruses

The viruses used in this study were derived from BAC-generated Wt TB40/E expressing GFP under the SV40 promoter (51). TB40/E recombinant viruses containing mutations in the miR-US5-2 and miR-US25-2 pre-miRNA sequences were generated by galactokines (galK)-mediated recombination (52). Briefly, the galK gene was inserted into the region of the miRNA hairpin using homologous recombination (miR-US5-2 galK F: CTCCATATATTTGTTACGATAGAATACGGAACGGAGGAGGCTTTCGCCACACCTATCCTGAAAGCGTTGCATTCTT CCTGTTGACAATTAATCATCGGCA; miR-US5-2 galK R: CGTATGCACAAA AGGTATGTGTGAATGGAAATACATGATGAATGTCACATCACGCAAAGCAGCCGTGGGAATGGTCTCAGCAAAAGTTCGATTTATTCAAC; dmiR-US25-2 galk F: AAATCCGAGAGGTAGCCAACAGCAGGGAGCCAGATGACGACGGACGTCGGGACCGACGGACGCGACTCGGGGTCCTcctgttgacaattaatcatcggca; dmiR-US25-2 galk R CCCCGCTTCGCCACCCCGCGCCCGCCCCAGTTTATACTCTCCGACGCCCCGTCCAACGCGCCTGTGGAGGGCCAATctcagcaaaagttcgattta). In the second recombination step, galK was removed using oligos that encompass the pre-miRNA sequence (miR-US5-2 oligo F: AGGCTTTCGCCACACCTATCCTGAAAGCGTTGCATTCTTTTCCTATCGTGTGACGATGTCTTTACCATTCCCACGGCTGCTTTGCGTGATGATGACATTC; miR-US5-2 oligo R: GAATGTCATCATCACGCAAAGCAGCCGTGGGAATGGTAAAGACATCGTCACACGATAGGAA; miR-US25-2 oligo F: CCAGTTTATACTCTCCGACGCCCCGTCCAACGCGCCTGTGGAGGGCCAATAGGACCCCGAGTCGCGTCCGTCGGTCCCGACGTCCGTCGCCATCTGGCTCAGAATGCAACGCTTTCAGGATAGGTGTGGCGAAAGCCT; miR-US25-2 oligo R: gagccagatggcgacggacgtcgggaccgacggacgcgactcggggtcctattggccctccacaggcgcgttggacggggcgtcggagagtataaactgg) or a Dvl3-targeting shRNA was designed (TGC TGT TGA CAG TGA GCG GCCCACTTTCTCCTACCAATA TAG TGA AGC CAC AGA TGT A TATTGGTAGGAGAAAGTGGGG TG CCT ACT GCC TCG GA) and amplified with the following sequences: (shRNA amp US25-2 F: AATCCGAGAGGTAGCCAACAGCAGGGAGCCAGATGGCGACGGACGTCGGGACCGACGGA CGCGACTCGGGGTAATT GGTATAT TGC TGT TGA CAG TGA GCG and shRNA amp US25-2 R: CGCTCCCCGCTTCGCCACCCCGCGCCCGCCCCAGTTTATACTCTCCGACGCCCCGTCCAACGCGCCTGTGGAGGGCCAAT GAA GTC CGA GGC AGT AGG CA). All viruses were propagated and titered on NHDF using standard techniques. To assess growth kinetics, NHDF were infected at an MOI of 3 for single-step growth curves or an MOI of 0.01 for multi-step growth curves for 2 hr. Cell-associated and supernatant virus was harvested at multiple time points post-infection and titered by plaque assay on NHDF.

### Reagents

The 3’ UTR of human Dvl3 was amplified by PCR from fibroblast genomic DNA and cloned downstream of the Renilla luciferase gene in the psiCHECK-2 dual luciferase reporter using the XhoI and NotI restriction sites and the following primers (Dvl3 F1: GCGG CTCGAG ccccagtgagttctttgtgg; Dvl2 R1: CGCCGGCGGCCCCTCACAACAACCCTGTGAGG). siGENOME RISC-free control siRNA (Neg; Dharmacon) and Dvl3 siRNA (s675; Thermofisher) were used in transfection experiments. Double-stranded miRNA mimics were custom designed and synthesized by Integrated DNA Technologies. The following commercial antibodies were used: Dvl3 (3218; Cell Signaling Technology), GAPDH (ab8245; Abcam) and HCMV IE2 (MAB810; Sigma Aldrich). Pen-N3 and Pen were purchased from Millipore Sigma. Wnt3a was purchased from R&D Biosystems. LiCl was purchased from Sigma.

### PAR CLIP and Bioinformatics analysis

293T cells stably transduced with pLCE-miR-US5-1/5-2 were expanded in log phase and pulsed with 100uM 4-thiouridine 2hr prior to addition of 5ng/mL of EGF. Cells were cultured for an additional 12 hours, washed with ice-cold phosphate buffered saline and cross-linked at UV 365 nm. To immunopurify Ago-bound RNAs, cell pellets were lysed in NP-40 lysis buffer and Ago complexes immunopurified with antibodies to pan-Ago (Diagenode) as previously described (53). cDNA libraries were sequences on the Illumina platform. Raw sequencing reads were obtained in FASTQ format. FASTQ files were concatenated, adapters removed and sequences collapsed with the FASTX-toolkit (http://hannonlab.cshl.edu/fastx_toolkit/), removing reads that contained an N or were <12 nt in length after adapter removal. Processed reads were aligned to the human genome (HG38) with Bowtie. Mapped reads were run through PARalyzer (29), requiring at least three reads per cluster. The PARalyzer distribution file was subsequently run through microMUMMIE (30) to predict additional Ago interaction sites. Ago interaction sites aligning to 3’ UTRs of human protein-coding transcripts were annotated for canonical miR-US5-1 and miR-US5-2 seed matches (>=6mer2-7).

### Luciferase assays

For dual luciferase assays, HEK293 cells were seeded into 96 well plates and transfected with 100ng of psiCHECK-2 vector and 100 fmol of negative control or miRNA mimic using Lipofectamine 2000 (Invitrogen). 24 hours post-transfection cells were harvested using the Dual-Glo Reporter Assay kit (Promega) according to the manufacturer’s instructions. For luciferase reporter assays, HEK293 cells were seeded into 96 well plates and transfected with 100ng of TOPflash (12456; Addgene) or FOPflash vector (12457; Addgene) as well as an 20ng Renilla luciferase-expressing control plasmid (SV40Rluc) and 100 fmol of negative control or miRNA mimic using Lipofectamine 2000. Cells were serum-starved for 16 hours and stimulated with 150ng/mL of Wnt3a or 25mM LiCl for 8 hours. Luminescence was detected using a Veritas microplate luminometer (Turner Biosystems), All experiments were performed in triplicate and presented as mean +/- standard deviation.

### Immunoblotting

Cells were harvested in protein lysis buffer (50mM Tris-HCl pH 8.0, 150mM NaCl, 1% NP40, and protease inhibitors), loading buffer (4X Laemmli Sample Buffer with 2-mercaptoethanol) was added, and lysates were incubated at 95°C for 5 min. Extracts were loaded onto 4–15% acrylamide gels (Biorad), transferred to Immobilon-P membranes (Millipore), and visualized with the specified antibodies. The relative intensity of bands detected by Western blotting was quantified using ImageJ software.

### Latency and reactivation assay

Differentiated hESCs were infected with the indicated viruses at an MOI of 2 for 48hr, or were left uninfected, in stem cell media (Iscove’s modified Dulbecco’s medium [IMDM] [Invitrogen] containing 10% BIT serum replacement [Stem Cell Technologies], penicillin/streptomycin, stem cell factor [SCF], FLT3 ligand [FLT3L], interleukin-3 [IL-3], interleukin-6 [IL-6] [all from PeproTech], 50uM 2-mercaptoethanol, and 20ng/ml low-density lipoproteins). Pure populations of viable, infected (GFP^+^) CD34^+^ HPCs were isolated by fluorescence-activated cell sorting (BD FACSAria equipped with 488-, 633-, and 405-nm lasers and running FACSDiva software) and used in latency assays as previously described (37, 38). Briefly, cells were cultured in transwells above irradiated stromal cells (M2-10B4 and S1/S1) for 12 days to establish latency. Virus was reactivated by coculture with NHDF in RPMI medium containing 20% FBS, 1% P/S/G, and 15ng/ml each of G-CSF and GM-CSF in an extreme limiting dilution assay (ELDA; (39)). GFP^+^ wells were scored 3 weeks postplating and the frequency of infectious centers was using ELDA software. In some experiments, 10uM Pen-N3 or Pen was added to the latency culture and replenished after 5 days.

### Quantitative PCR for viral genomes

DNA from CD34^+^ HPCs was extracted using the two-step TRIZOL (Thermofisher) method according to the manufacturer’s directions. Total DNA was analyzed in triplicate using TaqMan FastAdvanced PCR master mix (Applied Biosystems), and primer and probe for HCMV *UL141* and human β-globin as previously described (54). Copy number was quantified using a standard curve generated from purified HCMV BAC DNA and human β-globin-containing plasmid DNA, and data were normalized assuming two copies of β-globin per cell.

### Statistical analysis

Statistical analysis was performed using GraphPad Prism software (v10) for comparison between groups using student’s t-test, one-way or two-way analysis of variance (ANOVA) with Tukey’s post-hoc test or Bonferroni’s multiple comparison test as indicated. Values are expressed as mean +/- standard deviation or standard error of the mean, as indicated in the figure legends. Significance is highlighted with p<0.05.

## Conflict of Interest

The authors declare no competing financial or non-financial conflicts of interest

## Funding

This work was supported by the National Institutes of Health (P01 AI127335 to M.H.H. and P.C.). The funder had no role in study design, data collection and interpretation or the decision to submit the work for publication.

## Data Availability

The authors confirm that the main data supporting the findings in this study are available within the article and its supplemental material.

